# Pleiotropic effects drive correlation between body mass index and cortical myelination

**DOI:** 10.1101/274134

**Authors:** Lisa Ronan, Nenad Medic, Paul C Fletcher

## Abstract

**Background:** Epidemiological studies have reported significant associations between obesity and neurocognitive decline. Understanding these associations will require deeper analyses of how body mass index (BMI) and brain structure are related. Here we explore the extent to which shared genetic factors (pleiotropy) govern the association between BMI and cortical myelination.

**Methods:** Statistical models of bivariate heritability were applied to structural MR image data from a cohort of monozyogotic and dizygotic twins. Estimates of phenotypic and genetic correlation between BMI and cortical myelination were derived. A co-twin control design based on monozygotic twins was used to test the hypothesis of a causal relationship between BMI and myelination. The variation in the genetic correlation across the cortex was compared with the average statistical enrichment of genes associated with obesity derived from data from the Allen brain atlas.

**Results:** Statistically significant phenotypic and genetic correlation between BMI and cortical myelination was observed across the cortex. Taking the heritability of each trait into account, approximately 80% of the phenotypic correlation between the traits was accounted for by shared genetic factors.

Intra-pair differences between traits in monozygotic twins failed to support a causal relationship. Moreover, variation in genetic correlation across the cortex was significantly associated with the statistical enrichment of genes related to obesity.

**Conclusions:** These results support the hypothesis that pleiotropic effects drive the association between BMI and cortical myelination. This observation may help to explain the co-occurrence of obesity in neurocognitive decline and mental health disorders characterized by changes in myelination and oligodendrocyte function.

## 1. Introduction

Epidemiological studies have reported a significant link between obesity and neurocognitive decline and dementia risk (1-3). Relevant to this, increased body mass index (BMI) has been associated with a number of brain structure and functional changes (4). For example, from middle-age onwards, structural brain age is estimated to appear ten years older in overweight and obese individuals than their lean, age-matched counterparts (5). However the precise pathways linking BMI and brain are not well defined. Some studies suggest that changes in the brain may predate and possibly even lead to the onset of obesity. Indeed it has been observed that brain changes exist in subjects genetically predisposed to obesity before those subjects become obese (6). On the other hand, it is postulated that elevated BMI may precipitate a cascade of changes that influence the brain, either through endocrine dysfunction (3) or chronic inflammation (7,8). Although cytokines themselves are not typically neurotoxic (9), they may exacerbate neuronal damage in a number of ways (10). Conversely, severe caloric restriction or bariatric surgery have been demonstrated to provide some neuro-protective effects (11,12).

Here we explore the possibility that brain changes associated with obesity may be the result of pleiotropic effects, that is, the action of shared genes, rather than a causal relationship that has been speculated elsewhere. In a recent landmark study, it was demonstrated that genes related to obesity risk are significantly enriched in the CNS and involved in basic processes such as synaptic function and glutamate signaling (13). This raises the possibility that the association between BMI and brain structure may be due to shared genetic influences although this has not yet been established.

Twin data are often used to address such questions of pleiotropy. Monozygotic twins share 100% of their genome while dizygotic twins share just 50%. Structural equation models (SEM) make use of this fact (14), and can be constructed to assess heritability, which is a measure of how much of trait variance is attributable to genetic variance. In the bivariate case it is possible to assess the genetic correlation between traits, that is, the degree to which traits share genetic factors. In general a high genetic correlation may imply a significant phenotypic correlation although this is not necessarily the case given that the influence of genes one or both traits may be minimal. Thus, in order to assess the contribution of shared genetic influences to a phenotypic correlation, the square root of the heritability of each trait is multiplied by the genetic correlation between traits. Obesity itself is highly heritable (between 70 - 80% (15)) as are various brain parameters (16), suggesting that if a genetic correlation exists, it may explain the observed phenotypic correlation between traits. In turn this may have implications for our understanding of the biology relating BMI to structural and cognitive changes.

However the presence of a genetic correlation does not necessarily imply that two traits share the same genes. For example, if there is a causal relationship between the traits, the genetic factors that govern the first trait may in turn also influence the second (17,18). To distinguish between the action of shared genes and a causal association between traits, it is possible to use a co-twin control design to calculate the correlation between intra-pair differences of the traits in monozygotic twins (19). By controlling for the influence of the genotype in this way, a correlation between trait differences suggests a causal association between them.

Here, we capitalized on the rich data from the Human Connectome Project (20) to investigate the extent of pleiotropic effects between BMI, cortical myelination and cortical thickness. We used SEM models to estimate the heritability of traits, as well as their genetic correlation. We further estimated the percentage of phenotypic correlation observed between traits attributable to shared genetic influences. We used a co-twin control design to test the hypothesis of a causal relationship between BMI and cortical myelination given the latter’s significant role in learning, memory and cognitive health, and its association with a host of neurodegenerative diseases such as dementia and Alzheimer’s disease, as well as age-related cognitive decline (21). Finally we postulated that, if pleiotropic effects drive the phenotypic correlation between BMI and cortical structure, then the pattern of the association should correlate with the cortical expression pattern of genes associated with obesity. We used gene expression data from the Allen Brain Atlas (22) to test this hypothesis.

## 2. Methods and Materials

### 2.1 Subjects

Data from the Human Connectome Project (HCP) was used in this study (20). In total, imaging and demographic data from *145* twin pairs for whom genotyping was available were included. Genotyping was used to confirm zygosity of twins. The full dataset included 94 monozygotic and 51 dizygotic twin pairs. The mean age was 29 years (range 22 - 35 years). The mean BMI (defined as weight in Kg divided by height in meters squared) was 25*Kgm*^−2^ (range 19.7*Kgm*^−2^ - 44.7*Kgm*^−2^). There was no difference in the proportion of sexes across the twin groups (female to male: MZ 58:36, DZ 31:21),*X*^2^ = 0.03, p = 0.87.

### 2.2 MR Image Data

Structural imaging of this dataset has been reported elsewhere (23). Structural images were acquired on a 3T Siemens TIM Trio system employing a 32 channel head coil. A high resolution 3D T1-weighted structural image were acquired using a Magnetization Prepared Rapid Gradient Echo (MPRAGE) sequence with the following parameters: Repetition Time (TR) =2250 milleseconds; Echo Time (TE) =2.99 milliseconds; Inversion Time (TI)=900 milliseconds; flip angle =9^0^ degrees; field of view (FOV) = 256mm × 240mm × 192mm; voxel size =1mm isotropic; GRAPPA acceleration factor =2; acquisition time of 4 minutes and 32 seconds.

In order to verify image reconstruction quality and for purposes of calculating specific imaging parameters, we applied the FreeSurfer software (version 5.3) (24-26) to the HCP structurally preprocessed image files for each subject where available to generate surface reconstructions for each subject. Surface reconstruction processes were conducted in native space.

### 2.3 Cortical myelination and thickness

We chose to examine the pleiotropic effects between cortical myelination and cortical thickness based on previous studies which suggested linked between these traits and BMI (4,5). As previously reported (27), the myelin content of cortical areas co-varies with the intensities from T1w and T2w images. A surrogate of cortical myelin was generated by taking the ratio of the T1w to T2w signal intensity across the cortex. Cortical maps of this ratio were available for each subject in the HCP dataset. These maps were registered to the individual FreeSurfer cortical reconstructions for further analysis.

Measures of cortical thickness for each subject were derived in a standard way from the FreeSurfer surface reconstructions (28). Results from cortical thickness analysis were used to assess the degree of specificity of cortical myelination-based results.

### 2.4 Statistical Analysis

Structural equation modeling (SEM) was used to calculate the univariate heritability of BMI, cortical myelination. The heritability of a trait is defined as the proportion of phenotypic variance attributable to genetic variance (29). In the basic univariate genetic model, the ACE model, assumes that the variability in the observed variable can be explained by genetic and environmental factors, in which A represents the additive genetic factors, C the shared/common environmental factors and E unique/specific environmental factors. If statistically validated (i.e. no significant change in model fit as assessed by minus twice the log-likelihood value (−2LL) for the corresponding change in degrees of freedom (df), and lower values for the Akaike’s Information Criterion (AIC)), the ACE model may be simplified to an AE model in which trait-variance is a function of genetic and unique environmental factors only.

SEM methods were also used to calculated bivariate statistics such as the genetic correlation and the bivariate (shared) heritability between traits. Genetic correlation is a measure of the genetic overlap between traits. For example, a genetic correlation of 1 indicates that the genes of each trait overlap exactly. The phenotypic correlation between traits is a function of both genetic and environmental factors. Bivariate genetic models allow us to decompose phenoytpic correlation in to each of these factors to determine the extent to which phenotypic correlation is driven by genetic factors. The genetic contribution to phenotypic correlation is called the bivariate heritability which is a measure of the shared variance between traits, and is calculated as a product of the genetic correlation between traits and the square root of the heritability of each trait.

Bivariate genetic statistics were based on the Cholesky decomposition model. Data from this model was transformed in to the correlated factors model to generate estimates of genetic correlation (*r*_*g*_), and correlation due to shared and unique environmental factors (*r*_*e*_) (30) (see Figure S1). More detailed path calculations and are given in the supplementary material.

The statistical significance of the phenotypic, genetic and environmental correlations were estimated by dropping the relevant paths in the bivariate genetic models and testing model fit as assessed by the change in −2LL for the corresponding change in degrees of freedom. The correlation was deemed to be significant at alpha = 0.05 level. Further details are provided in the supplementary material.

We additionally sought to test whether there is a causal relationship between BMI and cortical myelination but calculating the correlation between the intra-pair differences in BMI and myelination (19). If there is a causal relationship, we expect that in a monozygotic twin pair, differences in BMI should equate to differences in myelin. This co-twin control design controls for the influences of the genotype on the relationship between BMI and myelin, thus if a relationship between the different in traits is observed in MZ twins it suggests a causal relationship between the traits.

Statistical analysis was carried out in R (version“Pumpkin Helmet”’) using the package “UMX”’ (http://github.com/tbates/umx, accessed August 2016) (31).

### 2.5 Cortical Maps

Applying complex models at a per vertex level across the cortex is problematic from the point of view of signal to noise, given the inherent variability of the cortex at an individual level, as well as the possibility that the signal of interest may manifest at a scale larger than a vertex. Applying models at larger levels also suffers from the problem of scale, as well as possible segmentation-bias, where results may differ depending on how a particular segmentation falls across the cortex. For this reason we applied we employed a random sampling technique to generate cortical parcellations with regions of different sizes in order to generate estimates of univariate and bivariate heritability unbiased by scale. Three distinct parcellation schemes were used to generate approximately 30, 60 and 100 regions in each scheme, with approximately the same surface area generated for each region per parcellation. Although randomly positioned, we further controlled for labeling bias by repeating our analysis ten times for each parcellation scheme. This resulted in 10×30, 10×60 and 10×100 different segmentation files for each hemisphere per individual (i.e. 30 random parcellation schemes per hemisphere per individual). Subsequent estimates of heritability and phenotypic and genetic correlation were based on these parcellations.

Each model (namely univariate and bivariate heritability) was calculated for each segment (i = 1:10) in each parcellation scheme (j = 30, 60, 100), generating associated path statistics for that model in that segment in that parcellation. From these we generated estimates of univariate heritability, phenotypic, genetic and unique environmental correlation for each segment in each parcellation. The final cortical map was generated by averaging the per vertex values of each statistic across the parcellation schemes;

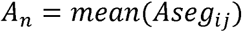

where *n* is the vertex index, *i* is the segment and *j* is the parcellation number (1:10). In doing this we produced a per vertex estimate of each statistic.

Multiple comparisons were controlled for using false discovery rate methods (32) using the p.adjust function in the statistics package of R.

Finally, in order to test if the relationship between BMI and cortical myelination was causal in nature, we calculated the correlation of intra-pair differences in BMI and cortical myelination in monozygotic twins for each parcellation.

### 2.6 Comparison with human cortical transcriptome

As a second stage to our analysis we sought to relate our identified pattern of genetic correlation across the cortex with gene expression data provided by the Allen Institute for Brain Science (22) using previously described techniques (33,34).

In brief, the Allen Human Brain Atlas (http://human.brain-map.org) is a publicly available online resource of microarray-based gene expression profiles for an anatomically comprehensive set of brain regions (22). The atlas is based on post-mortem tissue from 6 donors with no known history of neuropathological or neuropsychiatric disease, who also passed a set of serology, toxicology and RNA quality screens. The donors were a 24-year-old African American male, a 39-year-old African American male, a 57-year old Caucasian male, a 31-year old Caucasian male, a 49-year old Hispanic female, and a 55-year old Caucasian male.

The process of matching the microarray data from the AIBS to our MR image data has been described elsewhere (33,34). Briefly, the procedure involved segmenting our cortical data in to a series of 308 regions across both hemispheres. These regions correspond to regional gene expression profile.

Microarray data were averaged across all samples from all donors in the matching anatomical region across both hemispheres. The data were also averaged across probes corresponding to the same gene, excluding probes that were not matched to gene symbols in the AIBS data. Two MRI regions were excluded, because both the mean and the range of gene expression values in these regions were outliers compared with the other cortical regions of interest. The final output was a matrix of Z-scored expression values for each of 20,737 genes estimated in 306 MRI regions. In other words, for each cortical region, we generated a z-score value of the average gene expression for each of the 20,373 genes.

Using this data, we were able to calculate the average expression level of 248 genes previously linked to obesity risk (13) per region across the cortex and compared this map to the pattern of genetic correlation between traits. Analysis was restricted to the left hemisphere only as these genetic data were the most robust (22). A list of genes is provided in the supplementary material.

## 3 Results

### 3.1 Average values and univariate heritability

The mean BMI was 26*kgm*^−2^. The average cortical myelination varied across the cortex (left hemisphere 1.6, sd 0.12; right hemisphere 1.65, sd 0.15) (see Figure 1a). The heritability of BMI was 72% (p = 0). For statistically significant estimates of heritability of myelin the mean value in each hemisphere was 0.35 with range [0.2, 0.47] in the left hemisphere, and [0.22, 0.51] in the right hemisphere (see Figure 1b).

**Figure 1.**
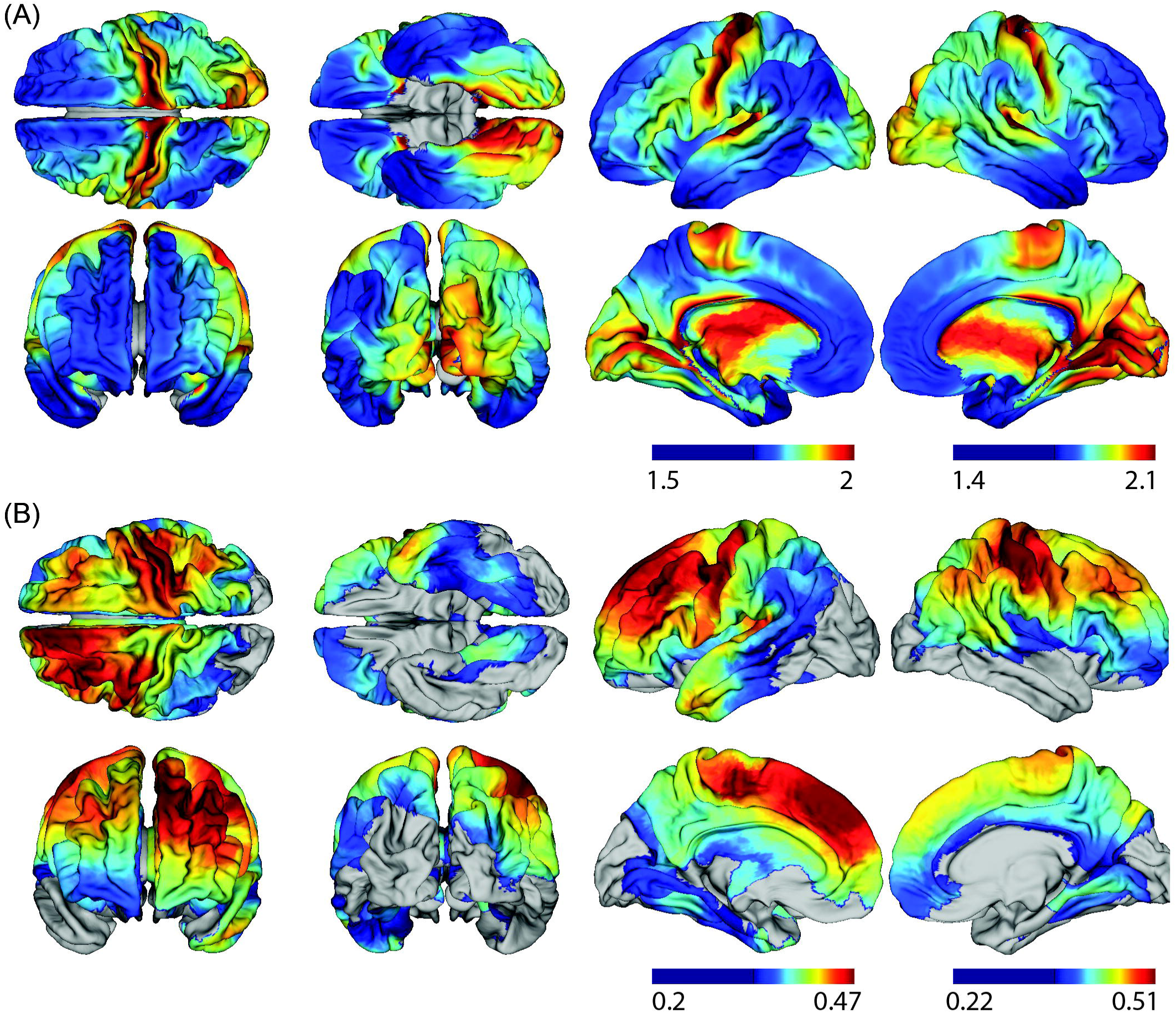
(A) Mean cortical myelination derived from the T1w-T2w ratio, and (B) heritability of cortical myelination thresholded for statistical significance.

### 3.2 Correlations between BMI and cortical myelination

Compared to the ACE model, there was a non-significant decline in fit for the AE model with the following average values across the cortex (−2LL = 1378; *df* = 8; AIC=235; *X*^2^ = 0.04, Δ*df* = 3; *p* = 0.99; ΔAIC = 5.9), indicating a non-significant effect of shared environment. Thus the AE model was chosen as the best-fit, most parsimonious model.

Across the cortex, almost all regions had statistically significant positive phenotypic correlations between BMI and myelination values (Figure 2a). The average value of the correlation coefficient was 0.24 (sd 0.04, average p-value 0.02) in the left hemisphere and 0.3 (sd 0.06, average p-value 0.02) although there was a considerable variation across the cortex and in particular a notable hemispheric asymmetry.

**Figure 2.**
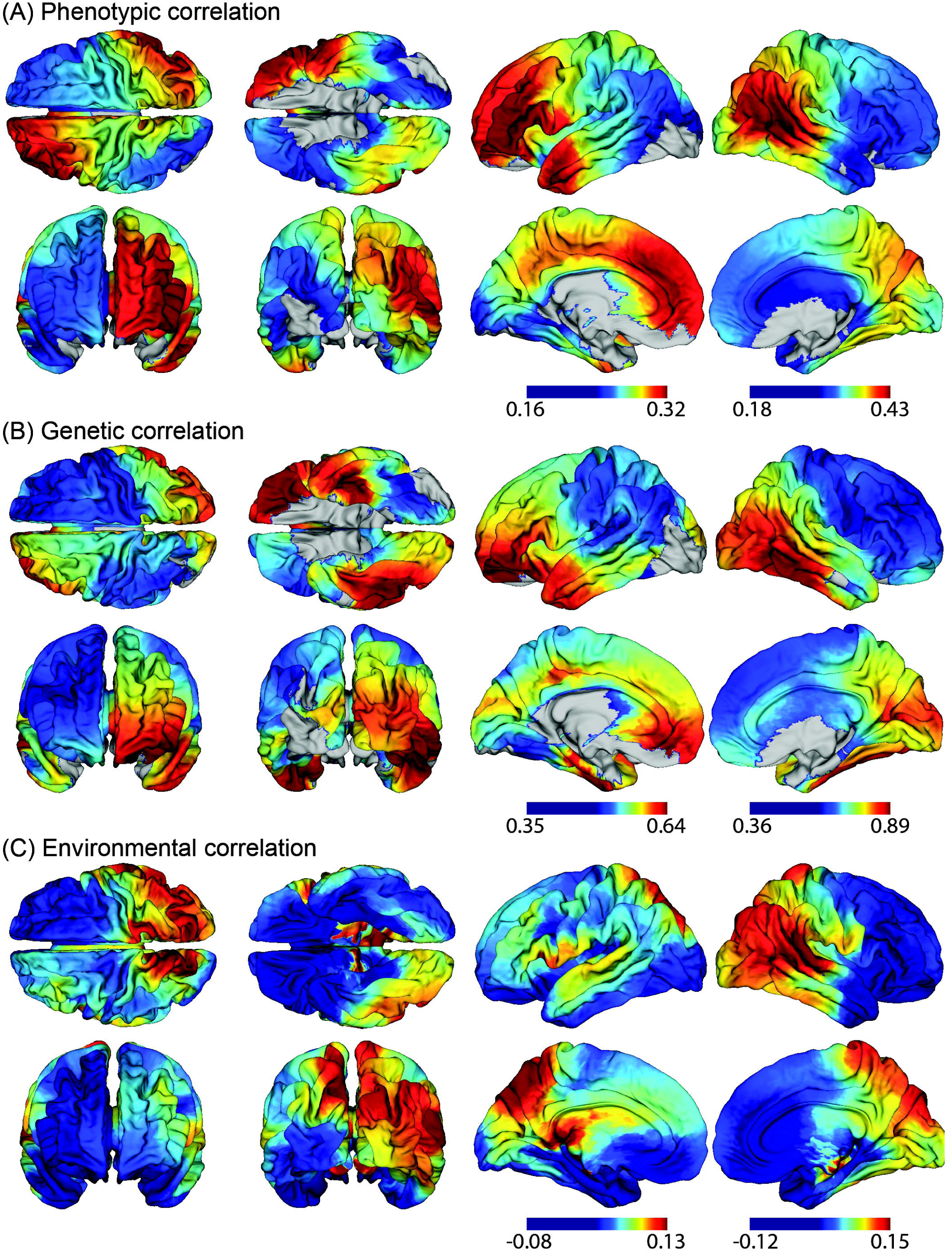
Cortical map of (a) phenotypic (b) genetic correlation and (c) unique environmental factors between BMI and myelination. Results for (a) and (b) are thresholded to illustrate only regions with statistically significant values. There were no statistically significant regions for unique environmental correlations.

Of note, this pattern of phenotypic correlation across the cortex was replicated for 358 unrelated individuals from the HCP dataset (with a similar age and sex distribution). See supplementary material for further details (Figure S2).

Genetic correlations were generally much higher across the cortex than the environmental correlations, suggesting that genetic factors were more significant in determining trait correlation. For regions of statistically significant genetic correlations (*r*_*g*_), estimates in the left hemisphere varied from [0.35, 0.64], with a mean of 0.47 (p = 0.03), while the right hemisphere varied from [0.36, 0.89] with a mean of 0.59 (p = 0.01)(Figure 2b). Squaring the genetic correlation gave the percentage of genetic influence shared between traits. In the left hemisphere this ranged from 12% - 40%, while in the right hemisphere it ranged from 13% - 79%.

The mean estimate of un-thresholded correlation due to unique environment (*r*_*e*_) was 0.04 (sd. 0.03, range [−0.08, 0.13]) in the left hemisphere, and 0.05 (sd. 0.05, range [−0.12, 0.15] in the right hemisphere (Figure 2c). After thresholding, there were no significant areas of correlation due to unique environmental factors.

When limited to regions of statistically significant genetic correlation only, the average proportion of phenotypic correlation attributable to genetic factors was 83% (sd 16%) in the left hemisphere and 76% (sd 23%) in the right hemisphere indicating that the phenotypic correlation was almost entirely influenced by pleiotropic factors (see Figure 3).

**Figure 3.**
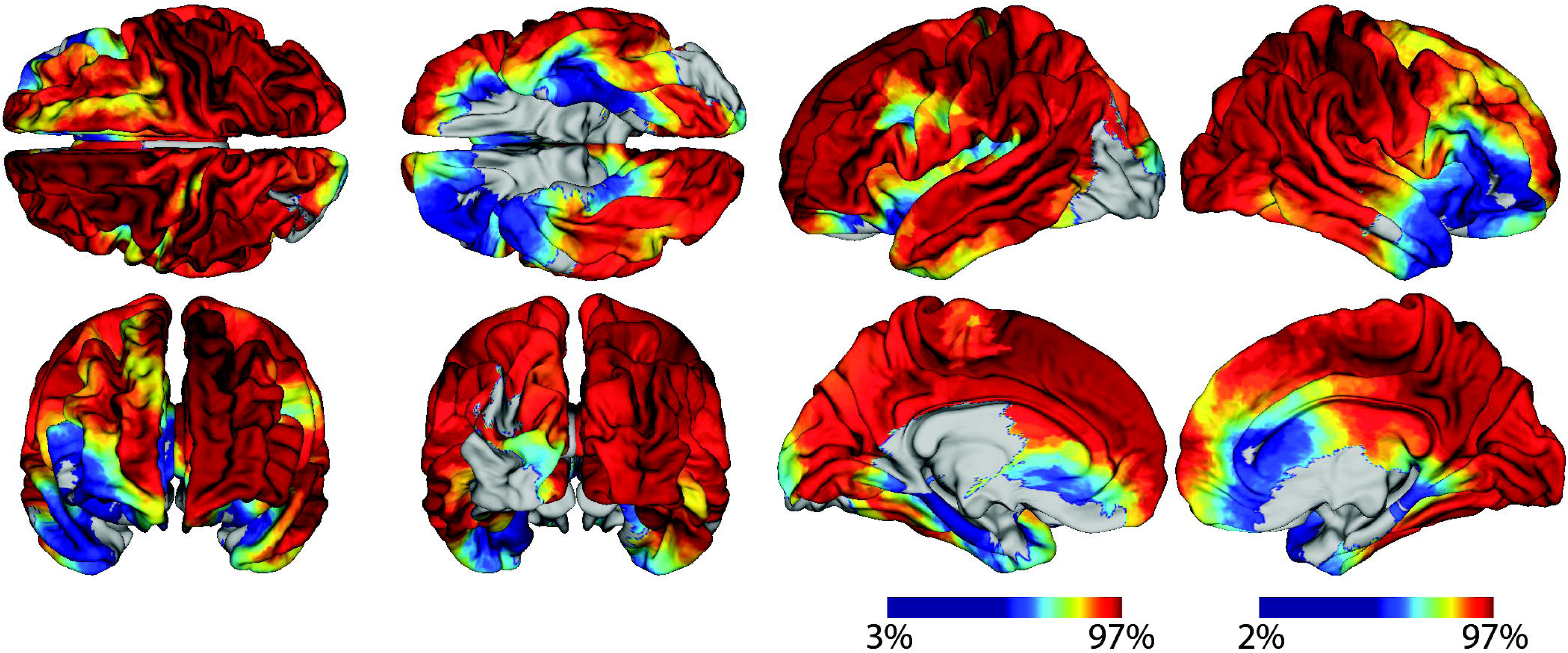
Map of the percentage of phenotypic correlation between BMI and cortical myelination attributable to genetic factors. This ratio is filtered for regions of statistically significant genetic correlation and is only calculable where all correlations are of the same sign.

### 3.3 Intra-pair differences in monozygotic twins

In a co-twin control design to test causality, the correlation between intra-pair differences in BMI and cortical myelination in monozygotic twins (n = 94) was calculated. There were no regions of the cortex where the correlation was statistically significant (see supplementary material Figure S3). These results fail to support the hypothesis that the association between BMI and cortical myelination is causal in nature.

### 3.4 Comparison with cortical transcriptome

There was a statistically significant correlation between the average statistical enrichment of obesity genes across the cortex and the pattern of genetic correlation between BMI and cortical myelination (R = −0.55 p < 0.0001) (Figure 4).

**Figure 4.**
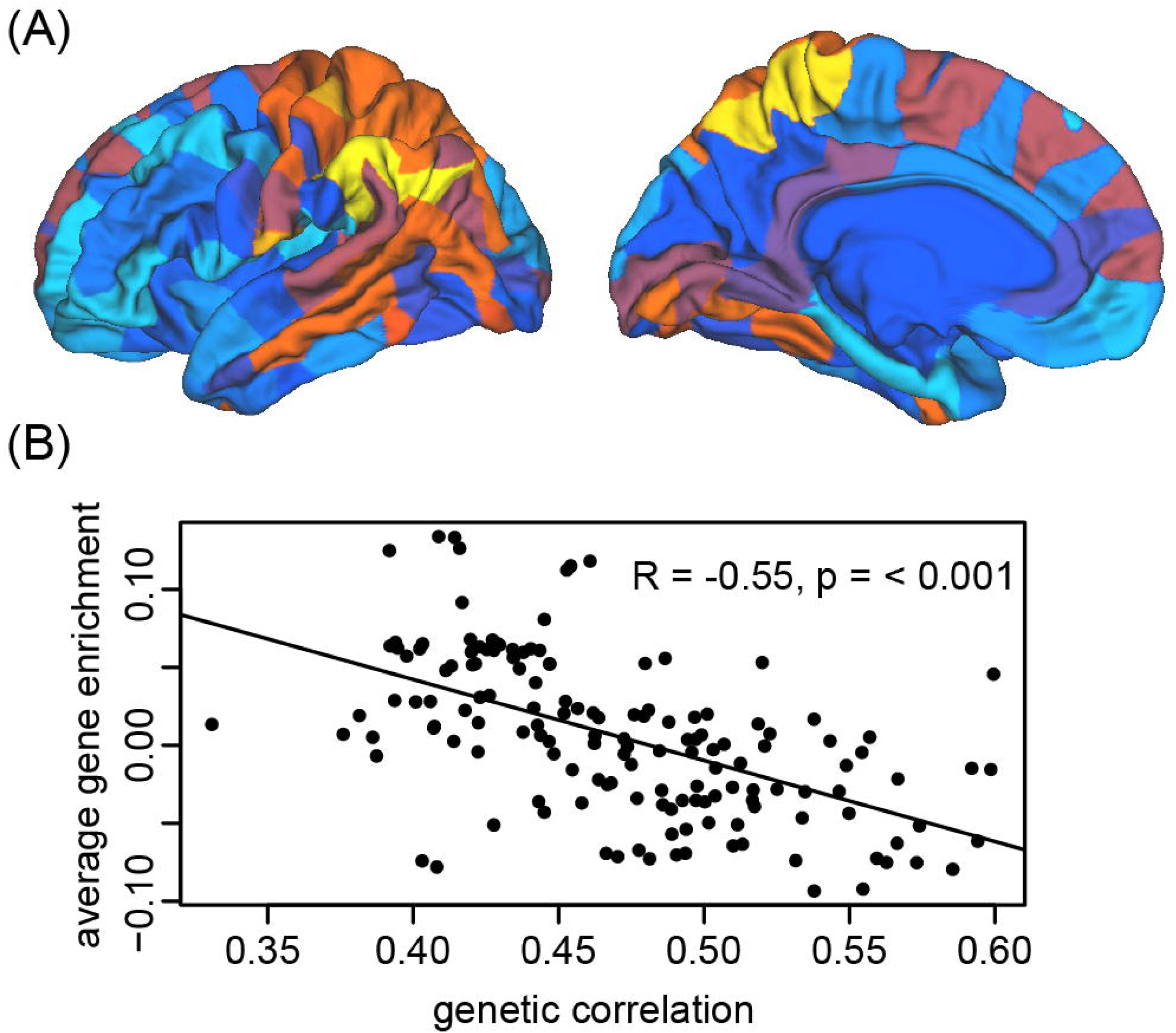
(a) Cortical map of average enrichment of genes related to obesity (Locke et al. 2015). (b) Correlation of average enrichment of genes related with obesity vs. genetic correlation between BMI and cortical myelination.

These results suggest that the pattern of genetic correlation between BMI and cortical myelination significantly co-varies with the expression patterns of genes related to obesity-risk.

### 3.5 Correlations between BMI and cortical thickness

There were no regions of statistically significant phenotypic or genetic correlation between BMI and cortical thickness.

## 4. Discussion

Our results demonstrate a significant genetic overlap between BMI and cortical myelination for large potions of the cortex. Taking the heritability of each trait in to account, we report that approximately 80% of the phenotypic correlation between BMI and cortical myelin was accounted for by shared genetic factors. The strength of phenotypic and genetic correlation between BMI and cortical myelination varied across the cortex, and it was, to a statistically significant degree, similar to the pattern of expression of genes related to obesity (13). Moreover, the relationship between intra-pair difference in BMI and cortical myelination in monozygotic twins failed to support a model of causal association between the traits. These results support the hypothesis that pleiotropic effects rather than a causal relationship govern the genetic association between BMI and cortical myelination.

The identification of a significant pleiotropy between cortical myelination and BMI may have important implications for our understanding of the biology underpinning the as yet unexplained association between BMI and cognitive decline and dementia risk. To date a large number of studies have related increased body mass in mid-life with an increased risk of cognitive decline and dementia in later years (2,35). Indeed it is suggested that an increased BMI in mid-life is associated with a two-fold higher risk of dementia on average (3,36). The results of the current study raise the possibility that genes associated with obesity have implications for cortical myelin which in turn may increase the rate of cognitive decline. Indeed, recent analysis has reported a significant genetic correlation between BMI and various aspects of cognition (37). More explicit studies of these associations are required to explore this hypothesis further.

In this analysis we failed to find any evidence of a significant genetic correlation between BMI and cortical thickness. These results suggest that our findings may be specific to cortical myelination as derived from the ratio or T1w to T2w image contrast (27). Other surrogates of cortical myelination such as magnetization transfer (38) may be used in future studies to confirm these results.

The influence of genes on brain structure has been demonstrated to vary with age. This raises the important caveat that the results of the current study must be interpreted for the age-range of the dataset (22-35 years). As such, the finding here that pleiotropic effects drive the phenotypic correlation between BMI and cortical myelination does not outrule the additional possibility that BMI may instigate a cascade of changes resulting in brain differences. This may be particularly relevant in middle-aged and elderly populations where longitudinal studies have demonstrated an increased risk of neurodegenerative disorders with increasing BMI (2,35).

Finally, as discussed elsewhere (17, 18), the presence of a genetic correlation fits not only with the possibility of the action of shared genes, but also with a causal model. However, the results of our monozygotic co-twin control experiment failed to support such a causal association. In a further exploration of our findings, we hypothesized that if a significant pleiotropy existed between BMI and cortical structure, then we would expect to see a correlation between the pattern of genetic correlation between traits and the pattern of expression of genes related to obesity risk. Our results confirmed this. However there are a number of important limitations to acknowledge in this approach to genetic enrichment analysis. In the first instance the results of the current study were drawn from a population of relatively young adults (mean 26 years). In contrast, gene express data from the Allen Institute was derived from older subjects (mean age 42.5 years). Given that recent work has demonstrated that age-related gene expression is genotype-dependent (39), it is therefore necessary to be cautious when interpreting the results of our gene enrichment analysis.

## Conclusion

There is a pervasive idea in the literature that obesity “causes”’ brain changes (40), and indeed there is good evidence that this might be the case. However these results and the results of other studies (13, 41) strongly indicate that structural brain changes observed in association with obesity may be the result of pleiotropic effects also. These findings may help explain the co-occurrence of obesity in cognitive decline and dementia-related illnesses characterized by changes in myelination and oligodendrocyte function.

## Acknowledgments

This work was supported by the Bernard Wolfe Health Neuroscience Fund and the Wellcome Trust (grant number RNAG/259). The authors would like to thank H Ziauddeen and D Gurdasani for their assistance in reading through the manuscript.

Data were provided [in part] by the Human Connectome Project, WU-Minn Consortium (Principal Investigators: David Van Essen and Kamil Ugurbil; 1U54MH091657) funded by the 16 NIH Institutes and Centers that support the NIH Blueprint for Neuroscience Research; and by the McDonnell Center for Systems Neuroscience at Washington University

